# Memory consolidation is linked to spindle-mediated information processing during sleep

**DOI:** 10.1101/206151

**Authors:** S.A. Cairney, A. Guttesen, N. El Marj, B.P. Staresina

**Affiliations:** Department of Psychology, University of York, UK; School of Psychology, University of Birmingham, UK

## Abstract

How are brief encounters transformed into lasting memories? Previous research has established the role of non-rapid eye movement (NREM) sleep, along with its electrophysiological signatures of slow oscillations (SOs) and spindles, for memory consolidation. More recently, experimental manipulations have demonstrated that NREM sleep provides a window of opportunity to selectively strengthen particular memory traces via the delivery of sensory cues. It has remained unclear, however, whether experimental memory cueing triggers the brain’s endogenous consolidation mechanisms (linked to SOs and/or spindles) and whether those mechanisms in turn mediate effective processing of the cue information. Here we devised a novel paradigm in which associative memories (adjective-object and adjective-scene pairs) were selectively cued during a post-learning nap, successfully stabilising next-day retention relative to non-cued memories. First, we found that compared to novel control adjectives, memory cues were accompanied by an increase in fast spindles coupled to SO up states. Critically, EEG pattern decodability of the associated memory category (object vs. scene) was temporally linked to cue-induced spindles and predicted next-day retrieval performance across participants. These results provide highly controlled empirical evidence for an information processing role of sleep spindles in service of memory consolidation.

## Introduction

Of the innumerable experiences we encounter throughout the day, only a fraction remain accessible to us in the future. Understanding the principles of consolidation, i.e., the mechanisms by which certain memory traces prevail across time, has remained a major challenge in memory research. Despite the boundaries of current understanding, there is general consensus that non-rapid eye movement (NREM) sleep plays an important role in memory consolidation, particularly for declarative memories that are initially reliant on the hippocampus. According to the *Active Systems* framework^1–4^, consolidation is driven by coordinated interactions between canonical electrophysiological signatures of NREM sleep. In concert with the depolarising up states of cortical slow oscillations (SOs, ˜.75 Hz), thalamocortical spindles (˜10-16 Hz) are thought to promote the stabilisation of memory traces in neocortical target areas. Accordingly, SOs and spindles have been robustly linked to overnight memory retention in a wide variety of studies^5–11^, signifying involvement of these oscillatory phenomena in sleep-dependent consolidation. Importantly, however, the functional role of SOs and spindles in memory processing during sleep has remained elusive.

Given the established role of sleep for memory, post-learning sleep periods might be used to strengthen memories even further by using experimental manipulations. Indeed, the recent advent of ‘targeted memory reactivation’ (TMR) paradigms has shown that NREM sleep provides a window of opportunity to bolster consolidation of specific memory traces^12–14^. In a typical TMR experiment, new memories are associated with auditory stimuli before a subset of the stimuli are replayed as memory cues during NREM sleep. For example, replaying environmental sounds associated with visuo-spatial memories improves the retrieval of cued relative to non-cued object-locations^15–18^. Likewise, replaying verbal foreign language cues facilitates the retrieval of cued (vs. non-cued) vocabulary translations^19, 20^. These findings are supplemented by a range of behavioural^21, 22^, neuropsychological^23^ and neuroimaging^24^ studies, indicating that TMR can be used to selectively strengthen declarative memories.

This begs the question of whether TMR in NREM sleep prompts selective memory benefits by exploiting the brain’s endogenous offline consolidation mechanisms. Of the neural oscillations unique to NREM sleep, spindles appear to be the prime candidate to support consolidation and task-related information processing in a targeted manner. Spindles tend to be more localised than more global SOs^25, 26^, and topographical differences in spindle activity have been observed following verbal vs. visuospatial learning^27, 28^. Moreover, recent work has linked spontaneous EEG activity in the spindle frequency band to memory processing in NREM sleep^29^, and BOLD increases in learning networks during NREM sleep are modulated by spindle parameters^30^. These observations suggest that spindles play a privileged role in mediating consolidation during sleep, and TMR may therefore bolster retention by generating finely-tuned windows of spindle-mediated information processing.

Here, we tested the prediction that experimental memory strengthening relies on deployment of sleep spindles and concomitant processing of task-related information. To this end, we devised a novel paradigm in which we controlled both the timing and content of memory consolidation. In particular, we presented sleeping participants with learning-related verbal cues and assessed (i) whether those cues triggered the emergence of SOs and/or spindles and (ii) whether those oscillatory events in turn mediate cue-specific information processing. Our findings critically revealed that valid memory cues – relative to novel control adjectives – led to an increase in spindle events modulated by SO up states. Moreover, during this time window of cue-induced spindle activity, the mnemonic association linked to the verbal cue could be reliably decoded, with the fidelity of this decoding predicting the behavioural consolidation benefits of TMR.

## Results

As shown in Figure 1, participants (n=46) encoded pairwise associations (adjective-object and adjective-scene pairs) before an initial test phase (T1), in which adjective recognition judgements (old or new) were made. Critically, for recognised adjectives, recall of the associated image (object or scene) was assessed. Half of the correctly recalled adjective-object and adjective-scene pairs were randomly assigned to a cued condition, such that the adjectives would be replayed during the offline period (TMR). The other half of the correctly recalled pairs were assigned to a non-cued condition. The to-be-cued adjectives were also intermixed with a number of non-encoded control adjectives. Immediately after T1, participants either took a 90-min nap (nap group; n=27) or remained awake for the same period of time (wake group; n=19). In the nap group, TMR took place during NREM stages N2 and N3. In the wake group, TMR coincided with a working memory task to prevent participants from directly attending to the cues^17, 20^. After the offline period, participants completed a second test (T2), before returning after an additional night of sleep to complete a final test phase (T3).

**Figure 1:**
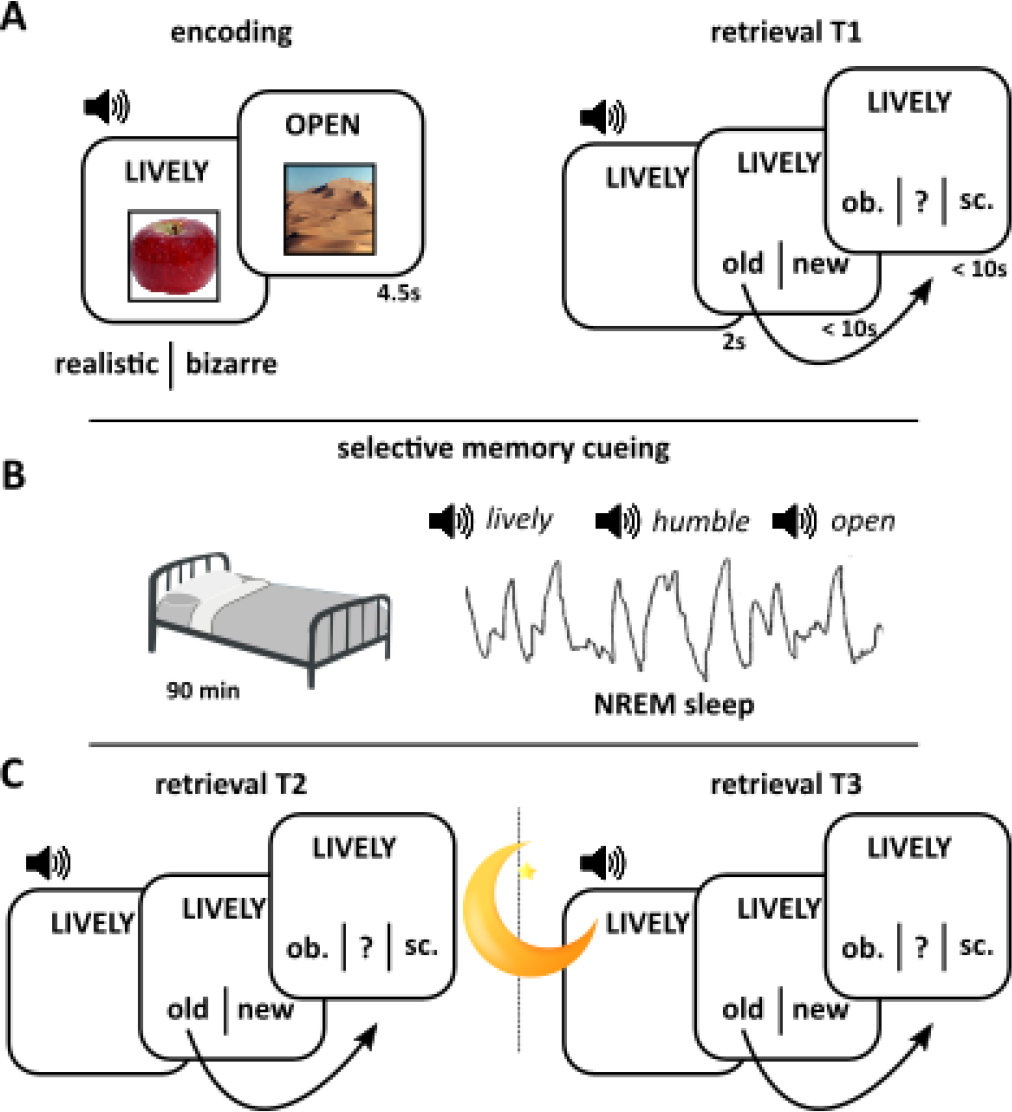
Paradigm. **A.** During encoding, participants were presented with 50 adjective-object and 50 adjective-scene combinations (randomly intermixed) and indicated whether they elicited a realistic or bizarre mental image. Prior to encoding, participants performed a familiarisation phase for both the adjectives and the images (see methods). Approximately 5 minutes after encoding, participants performed the first retrieval session (T1) in which all previously seen (old) adjectives were intermixed with 50 previously unseen (new) adjectives and participants indicated whether they thought the adjective was old or new. In case of an “old” response, they were asked whether they also remembered the associated image category (object or scene) or whether they did not remember the associated category (“?” response option). If they indicated “object” or “scene”, another screen appeared (not shown) in which participants could type in the exemplar if they remembered it or simply type in a “?” if they did not. Adjectives were presented visually and acoustically throughout. **B.** In the NAP group, participants were set up for polysomnographic recordings and were given the opportunity to sleep for 90 minutes. Once they entered deep NREM sleep, (i) half of the adjectives for which the object category was remembered at T1, (ii) half of the adjectives for which the scene category was remembered at T1 and (iii) a matched number of novel adjectives (controls) were continuously played via external speakers (targeted memory reactivation, TMR). In a matched WAKE group, participants started with 30 minutes of playing the online game Bubble Shooter, followed by 30 minutes of performing a 1-back working memory task during which TMR was applied, followed again by 30 minutes of playing Bubble Shooter. **C.** After the offline period (T2), participants performed the same test as in T1, but with a new set of 50 lure adjectives. Finally, after a night of sleep, participants returned the next morning (T3) for another retrieval session, again with 50 new lure adjectives.

**Table 1:**
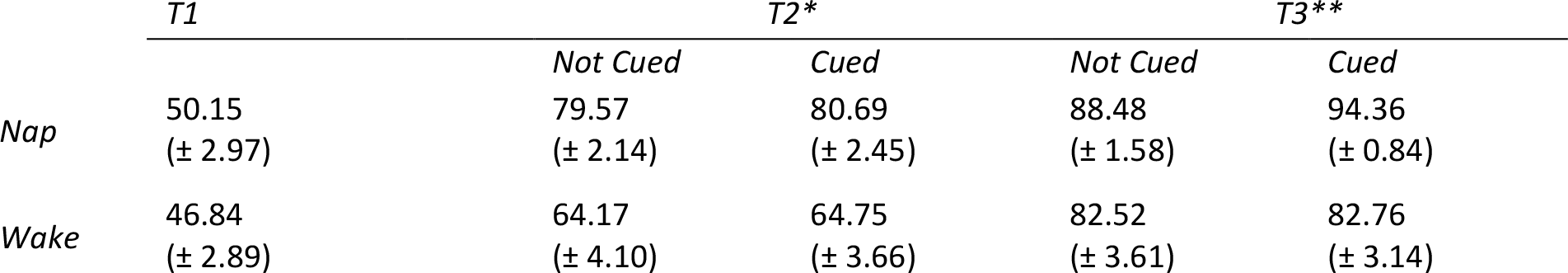
Category recall performance (%) at each test phase. Data are shown as means ± SEM. *Refers to the proportion of T1-recalled categories that were also recalled at T2. **Refers to the proportion of T2-recalled categories that were also recalled at T3.

|  | *T1* | *T2\** |  | *T3\*\** |  |
| --- | --- | --- | --- | --- | --- |
|  |  | *Not Cued* | *Cued* | *Not Cued* | *Cued* |
| *Nap* | 50.15<br>($\pm 2.97$) | 79.57<br>($\pm 2.14$) | 80.69<br>($\pm 2.45$) | 88.48<br>($\pm 1.58$) | 94.36<br>($\pm 0.84$) |
| *Wake* | 46.84<br>($\pm 2.89$) | 64.17<br>($\pm 4.10$) | 64.75<br>($\pm 3.66$) | 82.52<br>($\pm 3.61$) | 82.76<br>($\pm 3.14$) |

### Behaviour

Category recall at T1 did not differ between the nap and wake groups (*t*(44)=0.77, *P*=.45; see Table 1, Figure 2A). Category retention at T2 (i.e. the proportion of T1-recalled categories that were also recalled at T2) was significantly better after sleep than wakefulness (Group main effect: *F*(1,44)=17.10; *P*<.0001), but unaffected by cueing (TMR main effect: *F*(1,44)=0.19, *P*=.66; TMR*Group interaction: *F*(1,44)=0.02, *P*=.89) (Figure 2C). However, category retention at T3 (i.e. the proportion of T2-recalled categories that were also recalled at T3) revealed both a memory benefit of sleep (Group main effect: *F*(1,44)=9.34; *P*=.004) and a recall advantage from cueing (TMR main effect: *F*(1,44)=4.65, *P*=.04; TMR*Group interaction: *F*(1,44)=3.94, *P*=.05) (Figure 2D). The memory enhancing effects of TMR were driven by the nap group, who exhibited superior retention of cued relative to non-cued categories (*t*(26)=3.83, *P*=.001), whereas no such difference was observed in the wake group (*t*(18)=0.09, *P*=.93). Taken together, these results suggest that TMR in post-learning sleep augmented overnight consolidation processes, improving retention the following day. No differences were observed between object and scene recall (see supplementary materials).

**Figure 2:**
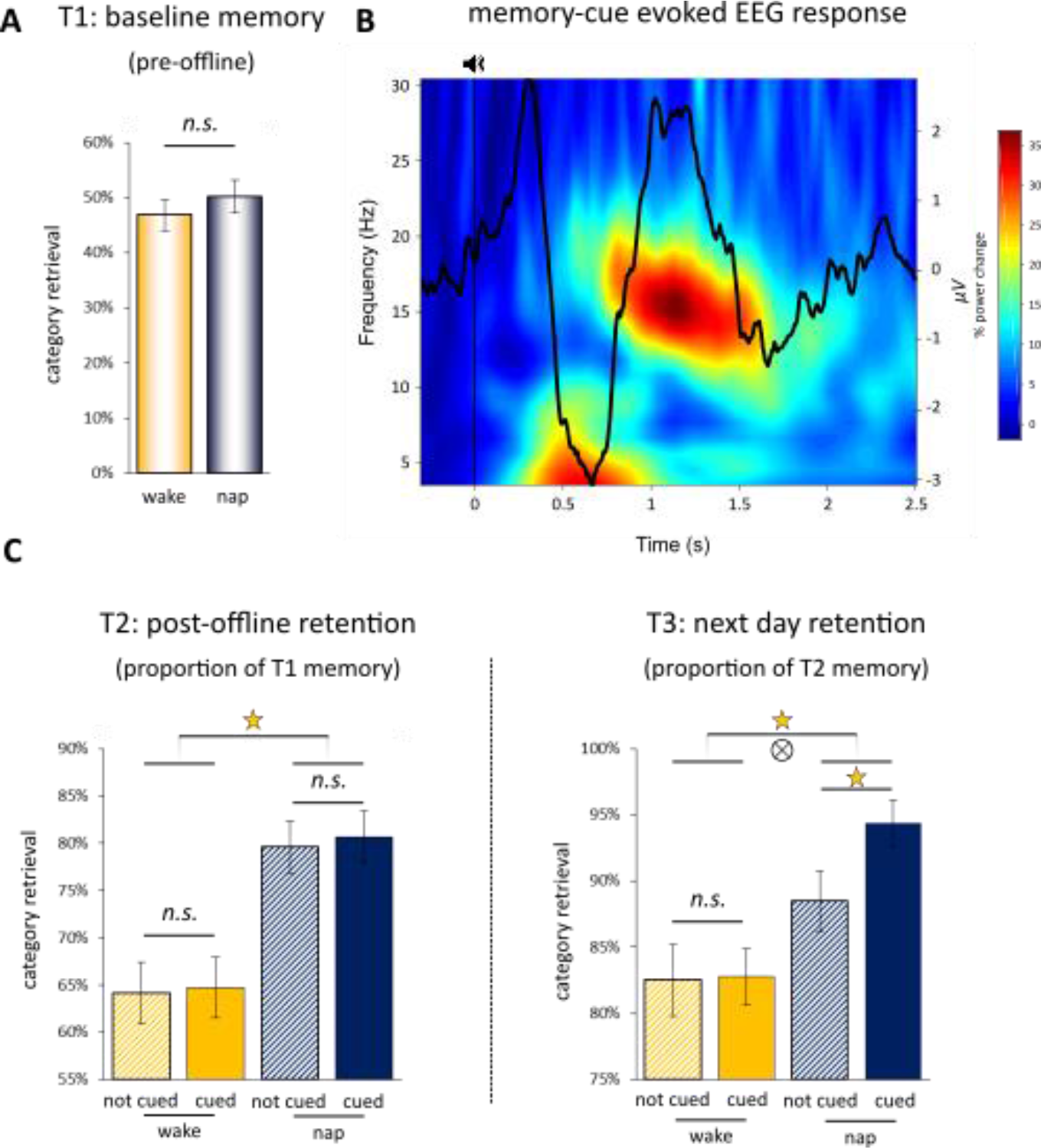
**A.** Behavioural results at T1 (pre-offline period). Bar graphs show mean (± SEM) accuracy for adjective-category retrieval for the NAP group (blue) and the WAKE group (orange). Note that 50% accuracy is not to be mistaken as chance performance given that participants had a “?” response option (see Figure 1A). **B.** Event-related potential (ERP) and time-frequency representation (TFR) evoked by the onset of memory cues. The figure depicts the unthresholded TFR along with the grand average ERP (both collapsed across all channels and then averaged across participants), revealing a strong increase of theta/slow spindle power in the evoked down state followed by an increase in fast spindle power in the ensuing up state. The statistically thresholded TFR map is shown in supplementary Figure S1. **C.** At T2 and T3, behavioural results are further separated into cued trials (solid fill) and not cued trials (hatched fill) and retrieval accuracy is expressed as proportions retained from the most recent memory assessment. Stars denote significant effects, ⊗ denotes an interaction effect.

### EEG

Sleep EEG data (i.e. time spent in each stage of sleep and spindle/SO event densities) are available in Table 2. Slow and fast spindle topography is illustrated in supplemental Figure S2.

**Table 2:**
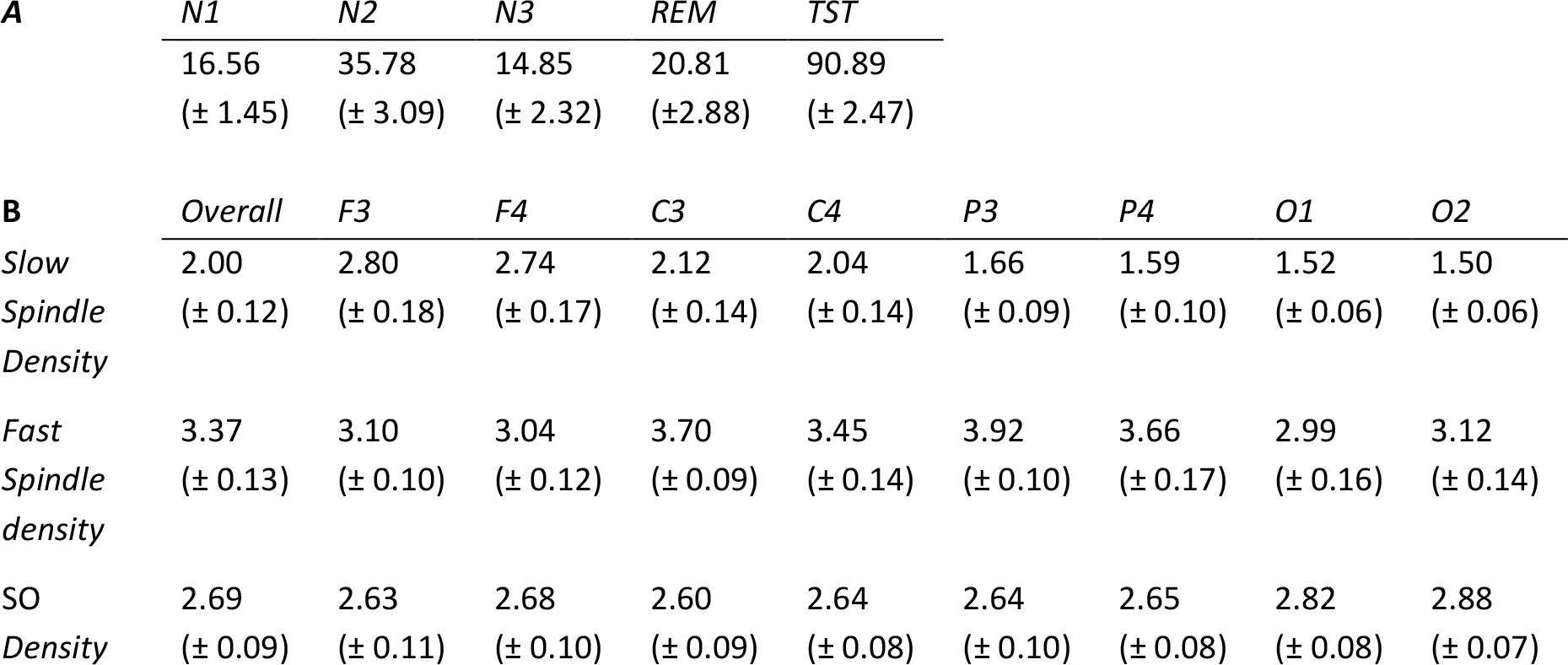
Sleep data. **A.** Time (min) spent in each stage of sleep and total sleep time (TST). **B.** Event density measures for slow spindles (10-13 Hz), fast spindles (13-16 Hz) and slow oscillations (SOs, 0.5-2 Hz) at each EEG channel. Density is calculated as the number of electrophysiological events divided by time spent in non-rapid eye movement (NREM) sleep stages N2 and N3. Data are shown as means ± SEM.

### Responses evoked by memory cues

As a first step, we explored the event-related potentials (ERPs) evoked by the auditory memory cues for previously presented (old) adjectives. As shown in Figure 2B and supplemental Figures S1 and S3, cues were followed by a pronounced ERP response resembling a SO/k-complex. Consistent with previous findings^19^, time-frequency representation (TFR) results showed that these cue-induced SOs harboured a theta/slow spindle burst in the down state which was followed by a fast spindle burst in the ensuing up state.

To more directly isolate the mechanisms engaged in the processing of mnemonic cues, we next asked whether evoked oscillatory responses might be able to distinguish between old cues and novel control adjectives. As shown in Figure 3A and 3B, the direct contrast revealed that old cues elicited a significant increase in oscillatory power in the fast spindle band (13-16 Hz) from ˜1.7-2.3 sec after cue onset (*P*<.05, corrected for multiple comparisons across channels, time and frequency). Topographical representation of the significant spindle increase revealed that the effect stemmed from left hemisphere electrodes, with a maximum at centroparietal sites C3/P3 (Figure 3C).

To fully characterise the observed spindle increase for old cues, we algorithmically detected discrete spindle events in C3 and P3 from 1-2.5 sec after the onset of old cues (see methods). Figure 3D displays the resulting grand average spindle across participants, aligned to the maximum of the detected spindle events. Not only does this result confirm that the oscillatory increase (Figure 3A and 3B) reflects discrete spindle events, but, as can be appreciated by the above-zero spindle centre, these spindles appeared to be modulated by the up states of concomitant SOs. To statistically quantify this observation, we derived the preferred phase of spindle-SO coupling for the detected events (see methods). Indeed, as shown in Figure 3D, the preferred phase of SO-spindle modulation clustered significantly around the SO up state (0°). This was confirmed when pooling all detected spindles across participants (Rayleigh test: z=35.8, *P*<1.4e-16; V test vs. 0: v=131.6, *P*<1.1e-16), as well as when deriving the preferred phases for each participant separately and then subjecting the resulting phase values to second-level statistics (mean preferred phase: 2°, Rayleigh test: z=8.3, *P*<1.3e-4; V test vs. 0: v=14.4, *P*<2.3e-05). In sum, these results show that old memory cues relative to new control adjectives elicit an increase in fast spindle activity which is localised to left hemisphere sites and modulated by the SO up state.

**Figure 3:**
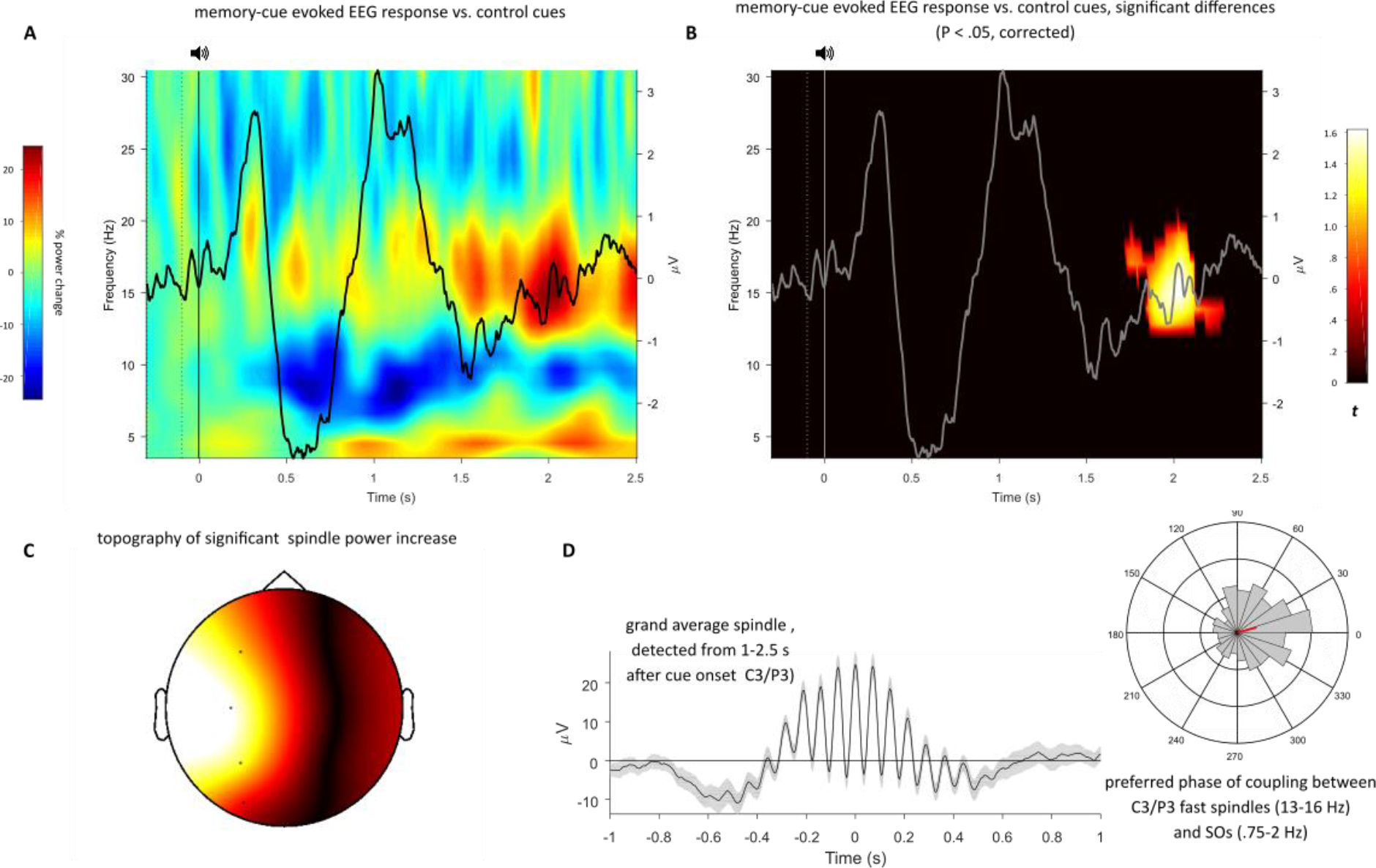
**A.** TFR difference map of responses elicited by old memory cues vs. new control adjectives, with the corresponding ERP for old cues superimposed. **B.** same as A, but after statistical thresholding (*P*<.05, corrected). Note the significant increase in fast spindle power (13-16 Hz) from ˜1.7-2.3 s post cue onset. **C.** Topography of the resulting cluster, revealing left-hemisphere specificity of the effect (see also supplemental Figure S4). **D.** *left*: Grand average (± SEM) of discrete spindle events detected from 1-2.5 s after onset of old memory cues at centroparietal sites. *right*: histogram of the corresponding SO-spindle modulation phases, pooled across all spindles at C3 and P3 across all participants (n=517, mean direction = 15°, shown in red).

### Information processing linked to fast spindles

If the surge of fast spindles was indeed indicative of cue-induced information processing, we reasoned that we should be able to decode from evoked EEG responses the categorical features (object vs. scene, Figure 1) of the images paired with the adjectives during learning. We used a representational similarity analysis (RSA) approach to tackle this question^31^. First, we derived a feature vector of 8 channels x 41 time points (spanning 200 ms at our sampling rate of 200 Hz) centred at each sample from -.2 s to 2.5 s relative to cue onset. Next, using Spearman correlations, we quantified both the within-category similarity (how similar is the EEG pattern of a given object-related adjective to the EEG pattern of all other object-related adjectives and how similar is the pattern of a given scene-related adjective to the pattern of all other scene-related adjectives) and the between-category similarity (how similar is the pattern of a given object-related adjective to the pattern of all scene-related adjectives and vice versa) at each time point. Category-specific information processing would be expressed as an increase in within-category similarity (collapsed across object- and scene-related adjectives) compared to the between-category similarity. As shown in Figure 4A, within-category similarity tended to exceed between-category similarity throughout the post-cue period. Importantly, however, the strongest effect that also reached statistical significance (*P*<.05, corrected for multiple comparisons across time) was observed from 1.76-2.12 s after cue onset, which overlaps precisely with the time window in which we observed the fast spindle increase for old relative to new cues (1.7-2.3 s, Figure 3B). Note also that converging evidence was obtained when using linear discriminant analysis (LDA, using k-fold cross-validation with k=10) to classify object- vs. scene-related cues, employing the same features and time windows as for the RSA analysis: the integrated classifier performance from 1.7-2.3 s (the window of fast spindle increase) showed a significant increase from chance performance (t(26)=2.3, *P*=.03 (twotailed)). However, the advantage of our RSA approach is that it provides, via the between-category similarity, a single measure that captures the pattern separability across objects and scenes (i.e. the smaller the between-category similarity, the more distinctive the object vs. scene patterns; see next step).

Lastly, we asked whether cue-specific information processing during the period of fast spindle increases bears any relevance for behavioural manifestations of consolidation. To this end, we assessed the correlation between (i) the relative benefit of cued vs. not cued items on T3 retention in each participant and (ii) the degree to which their evoked patterns of object- vs. scene-related adjectives are separable (1 - between-category similarity, averaged from 1.8-2.1 s after cue onset). As shown in Figure 4B, results revealed a significant positive relationship between the two variables (Pearson *r*=.45, *P*=.02; Spearman *r*=.41, *P*=.03). Taken together, these findings point to processing of learning-related information during the cue-evoked fast spindle surge, with the distinctiveness or fidelity of that processing predicting the behavioural benefit of TMR.

**Figure 4:**
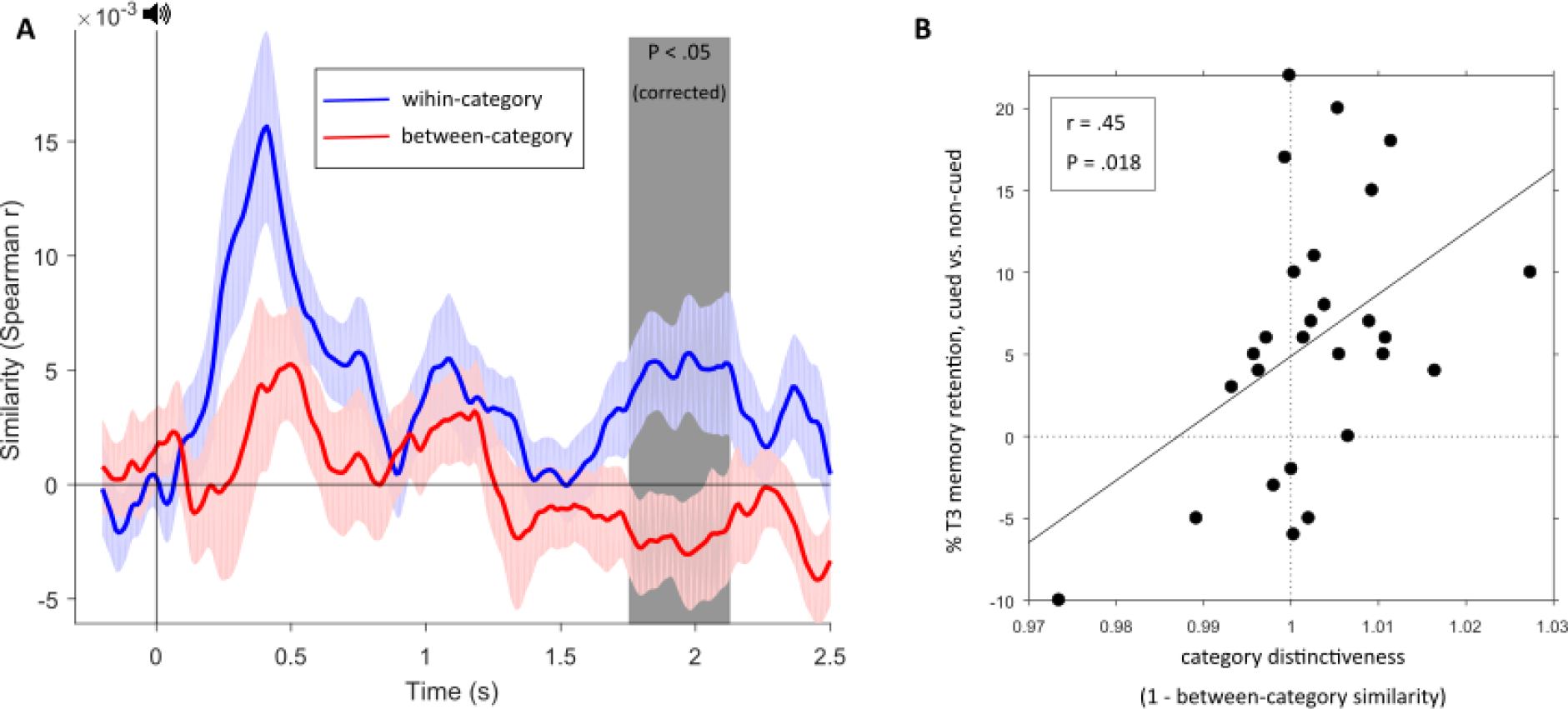
Information processing evoked by memory cues. **A.** Time course (mean ± SEM) of within- vs. between category similarity in response to old memory cues. Shaded area from 1.8-2.1 s highlights significant increase (P<.05, corrected for multiple comparisons across time). **B.** Correlation of category distinctiveness (1-between-category similarity from 1.8-2.1 s) and behavioural benefit of cueing on overnight consolidation across participants.

## Discussion

Our study reveals that spindle-mediated information processing is a central mechanism for offline consolidation, and that targeted memory reactivation (TMR) may exploit this mechanism to selectively strengthen cued memory traces. Specifically, we found that memory cues delivered in NREM sleep prompted a transient increase in spindle activity that was coupled with depolarising SO up states (Figure 3). During this surge in spindle activity, the categorical features of cued representations could be reliably decoded, with the extent of such decoding predicting the behavioural benefits of TMR (Figure 4).

The hierarchical nesting of spindles in SO up states is central to the *Active Systems* framework of sleep-dependent declarative memory consolidation^1–4^. Synchronised interactions between these oscillations are thought to be crucial for effective hippocampal-to-neocortical memory transfer. Interestingly, previous studies have indicated that fast spindles (13-16 Hz) in particular display strong temporal coupling with SO up states^6, 7, 25^. Corroborating a unique role for these oscillations in declarative memory processing, encoding of hippocampus-dependent memories has been found to enhance fast but not slow spindle activity in NREM sleep^32^, and fast relative to slow spindles are associated with greater BOLD activation in the hippocampus^33^. Here, auditory memory cues (but not novel control adjectives) elicited an increase in such fast spindle events, modulated by SO up states (Figure 3).

What is the functional significance of fast spindles for memory consolidation? Simultaneous EEG-fMRI recordings have shown that reactivation of learning networks is linked to spindle parameters during subsequent NREM sleep^30^. Moreover, olfactory memory cueing has been shown to evoke sleep spindles in task-relevant brain regions^34^. Although ceiling memory performance in these two studies precluded a direct link to consolidation, their findings suggest that a TMR-induced increase in fast spindle activity may reflect mnemonic processing in relevant hippocampal-neocortical networks. This view is substantiated by the left hemispheric specificity of the effect observed in our study (Figure 3C), which may reflect the verbal properties of the auditory cues.

More critically, during this transient, cue-induced increase in fast spindle activity, we were able to reliably decode the categorical features (i.e. object vs. scene) of the association linked to the verbal cue (Figure 4A), with the fidelity of this decoding predicting next-day retrieval performance across nap group participants (Figure 4B). One intriguing possibility is that spindles effectively gate activation towards category-specific cortical modules, leading to discriminable distributions of the spatiotemporal EEG patterns. Consistent with this notion, recent work applying EEG classifiers has shown that spectral power in the spindle range contributes to the ability to spontaneously decode previously learned materials in overnight recordings^29^. This converging evidence provides a good complement to the experimentally-evoked windows of spindle-mediated mnemonic activity observed in the current study. Finally, other work has shown that a short period following TMR-induced spindles is highly susceptible to disruption from additional auditory inputs, capable of abolishing the benefits of TMR for consolidation^19, 35^. Collectively, these findings point towards a crucial role of spindles in mediating information processing in service of sleep-dependent memory consolidation.

Mechanistically, modelling and empirical data suggest that spindle oscillations induce a massive Ca2+ influx into dendrites of pyramidal neurons, opening a molecular ‘gate’ for synaptic plasticity and, consequently, permanent network changes^36–38^. Finely-tuned windows of spindle activity, triggered by TMR, may therefore prime or ‘tag’ relevant synapses for plastic changes during subsequent periods of sleep. This may explain why the behavioural benefits of TMR observed in this study did not emerge until the following day, once the tagged representations had undergone additional overnight processing. Because sleep spindles are central to the proposed mechanism of synaptic priming via TMR, this may further explain why a memory benefit of cueing was not observed in the wake group (Figure 2). It should be noted though that the vast majority of studies employing nap-based TMR have reported improved retention of cued (vs. non-cued) memories immediately after sleep^15–17, 21^. However, our study is the first to integrate into a TMR paradigm the task of encoding arbitrary adjective-object or adjective-scene pairs. Thus, the spindle-mediated plastic processes induced by cueing in this context may have been insufficient to prompt a behavioural change without further overnight consolidation. Moreover, given recent views on the potential mechanistic overlap between online reactivation and offline consolidation^39^, another possibility is that T2 retrieval modulated interactions between prior TMR and subsequent overnight memory processing. How the memory effects of cueing are influenced by mnemonic activity in online and offline periods will be a fruitful avenue for future research.

Owing to the limited spatial resolution of scalp EEG monitoring, the putative role of hippocampally-generated ripples (>80 Hz oscillations) in our paradigm remains open. This is an important consideration as neuronal reactivations are mostly observed in conjunction with ripple events^40–42^, which are temporally nested within the oscillatory troughs of spindles^43–45^. The *Active Systems* framework postulates that these synchronised spindle-ripple interactions enable spindle oscillations to shuttle reactivated hippocampal representations to distributed neocortical sites during excitable SO up states^1–4^. Unifying our experimental paradigm with methods for detecting hippocampal ripples in the human brain (e.g. intracranial EEG) would thus provide exceptional insights into mnemonic processing in the sleeping brain. Notably, spindle-ripple interactions may reveal even greater detail on the informational content of decoded associations than spindle oscillations alone.

In sum, our findings suggest that experimental memory cueing generates finely-tuned windows of spindle-mediated information processing, which underpins the selective strengthening of cued representations. These findings not only offer mechanistic insights into the mnemonic impacts of TMR, but provide unique and highly controlled experimental evidence for the critical role of spindles in offline memory consolidation.

## Acknowledgements

This research was supported by a Wellcome Trust Institutional Strategic Support Fund (105624) through a University of York Centre for Chronic Diseases and Disorders (C2D2) Fellowship and a Medical Research Council (MRC) Career Development Award (MR/P020208/1) to SAC, and a Wellcome Trust/Royal Society Sir Henry Dale Fellowship to BPS (107672/Z/15/Z). The authors are grateful to Aidan Horner and Gareth Gaskell for fruitful discussions of the data.

## Methods

### Participants

A total of 83 participants took part in this study. However, 15 participants were excluded because they did not meet the performance criterion in the pre-sleep test (T1; see below). One further participant withdrew having not understood the necessary time commitments of the study. Of those participants remaining who took part in the nap version of the experiment, a further 21 were excluded for the following reasons: insufficient sleep such that at least one full round of targeted memory reactivation (TMR) could not be attained (9), exhibiting an arousal or awakening during TMR and not returning to non-rapid eye movement sleep stage N2/N3 (10) and computer malfunction (2). The analyses reported in this paper were thus carried out on 46 participants, who were assigned to a nap group (n=27, 20 female, mean ± SD age, 19.70 ± 1.51 years) or a wake group (n=19 females, mean ± SD age, 19.26 ± 1.15 years). Pre-study screening questionnaires indicated that participants had no history of sleep, psychiatric or neurological disorders, were not using any psychologically active medications. Participants were informed that they were taking part in a memory study, but were unaware that targeted memory reactivation (TMR) would take place. Written informed consent was obtained from all participants in line with the Research Ethics Committees of the Department of Psychology, University of York and the School of Psychology, University of Birmingham.

### Stimuli

#### Adjectives

250 adjectives were randomly selected from a longer list of 355^1^ for each participant. Mean (± SD) adjective length was 6.85 ± 1.84 characters and number of syllables ranged from 1-4. All adjectives were recorded in a female voice. Mean (± SD) duration of all included adjectives was 704 ± 146 ms.

#### Images: Objects and Scenes

100 images (50 objects and 50 scenes) were randomly selected from a set of 200^2, 3^ for each participant. Objects were images of everyday items, animals or food presented on a plain white background (e.g. an apple). Scenes were images of landscapes or places (e.g. a bowling alley). Care was taken to ensure that scenes contained sufficient background detail to be easily distinguishable from objects. The distinction between objects and scenes was clearly explained to participants.

### Procedure

Participants completed a short practice version (10 trials) of each experimental tasks to ensure that they fully understood the instructions. All responses were made via keyboard press on a PC. Experimental stimuli were always presented in random order.

#### Familiarisation

A familiarisation phase at the beginning of the experiment was designed to facilitate learning of the adjective-image pairs in the main encoding session. First, participants completed an adjective familiarisation task. On each trial, one of 100 adjectives (e.g. “exotic”) was presented aurally and displayed for 2.5 s on the computer screen. Participants indicated whether they considered the adjective to be emotionally positive or negative. Each trial was separated by an inter-stimulus interval (ISI) with a fixation cross for 1.5 s (± 100 ms random jitter). Next, participants completed an object/scene categorisation task. On each trial, one of 50 objects (e.g. butterfly) or 50 scenes (e.g. golf course) was displayed for 2.5 s. Participants indicated whether they considered the image to be an object or a scene (ISI=1.5s ± 100 ms).

#### Encoding

Participants learned randomised pair-wise associations between the adjectives and images presented in the familiarisation phase (100 adjective-image pairs). On each trial, a randomly selected adjective (e.g. “exotic”) was presented aurally and displayed above an object or scene (e.g. object: butterfly) for 4.5 s. To facilitate learning, participants were instructed to form a vivid mental image or story that closely linked the adjective and the object/scene, and then indicate whether the image they had formed was realistic or bizarre (ISI=1.5s ± 100 ms). For example, the mental image corresponding to the adjective “exotic” and the object butterfly would presumably be realistic as butterflies can be exotic creatures. Participants were informed that their memory for adjective-image pairs would be tested immediately afterwards.

#### Immediate Test (T1)

T1 included the 100 adjectives from encoding intermixed with 50 new adjectives that participants had not seen before (foils). On each trial of the test phase, an adjective (e.g. “exotic”) was presented aurally and displayed for 2 s. Afterwards, participants were asked to indicate whether the adjective was ‘old’ (i.e. they recognised it from the encoding phase) or ‘new’ (i.e. it was not seen at encoding) within 10 s. When participants provided a “new” response, they immediately moved on to the next trial (ISI=1.5 s ± 100 ms). When an “old” response was provided, participants were required to indicate whether the associated image was an object, a scene, or unknown to them. In order to ensure that object or scene responses were not mere guesses, participants also provided a typed description of the image they had remembered. Across all T1 trials where the category was correctly recalled, participants were able to correctly describe the image on the majority of occasions (mean ± SD: 80.95 ± 14.59%), demonstrating that the category responses reflected veridical memory.

#### TMR Set Up

Of the adjective-image pairs that were correctly recalled at T1 (i.e. when the adjective was correctly recognised and the associated image category correctly recalled), half were randomly allocated to the cued condition (i.e. for TMR), whereas the other half were assigned to the non-cued condition. This ensured that baseline category recall performance was balanced between the cued and non-cued memories. For example, if a participant correctly recalled 40 pairs at T1, then 20 of these would be assigned to the cued condition and the other 20 assigned to the non-cued condition. On occasions where there were an odd number of correctly recalled pairs, the spare item was randomly allocated to the cued or non-cued condition. To ensure that a sufficient number of stimuli were available for TMR in sleep, participants were required to correctly recall at least 14 objects and 14 scenes at T1. Participants that did not meet this criterion were excluded (n=15). The adjectives from pairs assigned to the cued condition were replayed during the TMR phase.

Importantly, an additional set of control adjectives that participants had not encountered at either encoding or T1 were randomly intermixed with the TMR stimuli. The number of control adjectives was equal to half the number of stimuli in the cued condition. For example, if there were 40 adjectives associated with correctly recalled categories in the cued condition, then a further 20 control adjectives would be added to the TMR set (total=60). Inclusion of the control adjectives enabled a direct comparison of brain activity during cued memory reactivation and non-specific, matched auditory stimulation.

#### Offline Period (Nap or Wakefulness)

The offline period began at ˜2pm and lasted 90 min. Participants in the nap group were left to sleep in a laboratory bedroom while their brain activity was monitored with polysomnography (set up before the study began). TMR was initiated when participants were in late NREM stage N2/early stage N3. The TMR set was presented in a randomised order (ISI=4s ± 200 ms) at a sound intensity of ˜40dB. After each full round of cueing, the adjectives were reshuffled and immediately presented again. Cueing continued for as long as participants were in sleep stage N2/N3, but immediately paused if they showed signs of micro-arousal or awakening, or moved into sleep stage N1 or rapid eye movement (REM) sleep. The cues were continued if participants re-entered sleep stage N2/N3 after an arousal.

Participants in the wake group played an online game (Bubble Shooter) for the first 30 min of the offline period. For the next 30 min, the TMR cues were presented continuously while participants completed a 1-back working memory task. This approach reduced the probability that participants directly attended to the cues during TMR^4, 5^. During the 1-back task, a series of random numbers between 0 and 10 were presented one after another in the centre of the screen. The task was to indicate whether the current number was the same as or different to the number one digit prior. After completing the 1-back task, participants played Bubble Shooter again for the remaining 30 min of the offline period. The mean (± SD) number of presentations per cue was 9.16 ± 3.67 in the wake group and 5.96 ± 5.85 in the sleep group.

#### Follow-Up Tests (T2 and T3)

Participants returned 6 hours later for a follow-up test (T2). This followed the same procedures as T1 with the single exception that new foil adjectives were used. The next morning (after a night of sleep), participants completed another test (T3). Again, this followed the same procedures as T1 and T2, but with a new set of foils.

#### Discrimination Task

After completing T3, participants were informed of the true purpose of the study and asked if they had been aware of the auditory cues in the offline period. To assess their explicit knowledge of the cues, participants were asked to complete an adjective discrimination task. On each trial, one of 100 adjectives from the encoding phase was presented aurally and displayed for 10 s. Participants were asked to indicate whether or not the adjective had been replayed during the offline period. Discrimination task data are available in the supplementary materials.

### Equipment

#### Experimental Tasks and TMR

All experimental tasks and TMR algorithms were implemented on a PC with MATLAB 2015a and Psychtoolbox version 3.0.13. In the wake group, adjective cues were presented via speakers connected to the task PC. In the nap group, cues were presented via a speaker mounted ˜1.5m above the bed, which was connected to an amplifier in a separate control room.

#### Polysomnography

An Embla N7000 PSG system with RemLogic version 3.4 software was used to monitor sleep. After the scalp was cleaned with NuPrep exfoliating agent (Weave and Company), gold plated electrodes were attached using EC2 electrode cream (Grass Technologies). EEG scalp electrodes were attached according to the international 10-20 system at 8 locations: frontal (F3, F4), central (C3, C4), parietal (P3, P4) and occipital (O1, O2), and each was referenced to an electrode on the contralateral mastoid (A1 or A2). Left and right electrooculography electrodes were attached, as were electromyography electrodes at the mentalis and submentalis bilaterally, and a ground electrode was attached to the forehead. Each electrode had a connection impedance of < 5 kΩ and all signals were digitally sampled at 200 Hz. Sleep scoring was carried out in accordance with the criteria of the American Academy of Sleep Medicine^6^.

### Data Analysis

#### Behaviour

Category recall was our primary measure of memory accuracy. We calculated for each participant: 1) the proportion of categories recalled at T1 that were subsequently recalled at T2, and 2) the proportion of categories recalled at T2 that were subsequently recalled at T3 (i.e. following a night of sleep). To avoid any ambiguity related to category memory, we excluded from our analyses any item that was incorrectly classified during the object/scene categorisation task. Across all participants, we excluded 162 items out of a possible 4600 (3.52%). Category recall scores were subjected to a 2 (TMR: Cued/Not-Cued) X 2 (Group: Nap/Wake) mixed analysis of variance (ANOVA).

#### EEG

EEG data were analysed with MATLAB (MathWorks), using the FieldTrip^7^ and CircStat^8^ toolboxes. The continuous sleep data were segmented into epochs from −1 s to 3 s around cue onset and subjected to a two-step artifact rejection procedure. In the first step, artifacts were automatically detected and removed based on the median plus/minus 3.5 inter-quartile ranges of both signal amplitude and gradients (the difference between two adjacent samples) of all epochs. In the second step, the remaining epochs were manually screened via FieldTrip’s visual summary functions and epochs containing amplitude, variance or kurtosis outliers were additionally removed. For TMR-locked analysis of event-related potentials (ERPs), data were high-pass filtered at 0.5 Hz and baseline-corrected with respect to the −200 ms to 0 ms window before cue onset. For time-frequency representations (TFRs), data were convolved with a 5-cycles hanning taper and spectral power was obtained from 4-30 Hz in 0.5 Hz frequency steps and 5 ms time steps. For analyses, participant-specific TFRs were converted into percent power change relative to a −300 ms to −100 ms pre-cue window.

For representational similarity analysis (RSA) of within- vs. between-category processing, a sliding window of 200 ms (in steps of 10 ms) was used to obtain, for each trial, a series of 8-channel-by-41-timepoints (200 Hz/5 ms sampling rate) EEG feature vectors. Using these feature vectors, Spearman correlations were then used to quantify, for each time point, the representational similarity across all pairwise combinations of trials. Next, we collapsed similarity values to capture (i) the average similarity of an object-related cue with all other object-related cues (i.e., within-object similarity), (ii) the average similarity of a scene-related cue with all other scene-related cues (i.e., within-scene similarity) and (iii) the average similarity of all object-related cues with all scene-related cues (i.e., between-object-scene similarity). Category-specific information processing was quantified as the difference between within-category similarity ((i) and (ii) combined) and between-category similarity (iii). Finally, the measure “1 - between-category similarity” was used to index the distinctiveness of category-specific information processing per participant (the greater the between-category similarity, the smaller the discriminability of object-vs. scene-related information).

All ERP, TFR and RSA analyses were corrected for multiple comparisons using FieldTrip’s cluster-based permutation method (1000 randomisations), including channel x time (ERP), channel x time x frequency (TFR) and time (RSA) as cluster-defining features.

#### Event Detection

Sleep data were partitioned according to the time (minutes) spent in each stage of sleep (N1, N2, N3 and REM sleep). Data scored as N2 or N3 were extracted from all EEG channels for spindle and slow oscillation analysis. For spindles, data were first bandpass filtered from 10-13 Hz (slow spindles) or 13-16 Hz (fast spindles) using a 4th order two-pass Butterworth filter. Next, we took the envelope of the resulting signal and determined an amplitude threshold as mean + 1.25 SD. A spindle was then defined as an event that surpassed that threshold for a minimum of 0.5 s and a maximum of 3 s. For SO detection, data were filtered from 0.8-2 Hz using a 4th order two-pass Butterworth filter. Next, zeros crossings were detected in the resulting signal and events with two successive positive-to-negative crossings spanning 0.8-2 s were taken forward to the next step. Here, the resulting candidate events’ trough and trough-to-peak amplitude were calculated and events surpassing mean + 1.25 SD of both these metrics were considered SOs. In both event detection procedures, automatically detected artifact samples (see above) were padded for ± 1 s and those samples were excluded prior to event detection.

For determining the preferred phase of SO-spindle modulation, we first identified spindles whose maximum occurred from 1-2.5 sec after onset of old memory cues (encompassing the interval in which we observed the spindle increase, Figure 3B). We then extracted a ± 1.5 s raw data segment around the spindle maximum (accommodating the maximum spindle duration of 3 s) and created one signal by filtering the data between 0.5 and 2 Hz and another signal by filtering the data between 13 and 16 Hz. For the lower frequency signal, instantaneous phase was extracted via the Hilbert transform. For the higher frequency signal, phase of the power envelope was extracted, again using the Hilbert transform. For each sample (601 samples, i.e. 3 s at 200 Hz sampling rate), the circular distance between the two phase time series was calculated and the mean resulting angle determined. For instance, if the spindle amplitude were to systematically peak at the SO down state (trough), the mean angle of the phase differences would be 180°. Conversely, if the spindle amplitude was to – as hypothesised – systematically peak at the SO up state (peak), the mean angle of the phase differences would be 0° (see also^9, 10^).

## Supplementary Materials

### Adjective Recognition

In each test, participants made recognition judgements (old or new) for adjectives before recalling the associated image category. The sensitivity index (*d’*) [Normalized (hits/(hits + misses)) – Normalized (false alarms/(false alarms + correct rejections))] was calculated for adjective recognition memory. We adopted a log-linear approach to safeguard this analysis against errors arising from 0 and 1 values: for each participant, 0.5 was added to the total hits and total false alarms, and 1 was added to the total signal (old) trials and total noise (new) trials^1^. Adjective recognition scores (*d’*) were applied to a 3 (Test: T1/T2/T3) X 2 (Group: Nap/Wake) mixed ANOVA. Recognition performance initially declined between T1 and T2, but then stabilised between T2 and T3 (Test main effect [Huynh-Feldt corrected]: *F*(1.78,88)=9.00, *P*=.001). However, these changes in performance were not modulated by the type of offline activity (Group*Session interaction [Huynh-Feldt corrected]: *F*(1.78,88)=2.36, *P*=.11). There was also no main effect of Group (*F*(1,44)=0.53, *P*=.47), indicating that overall adjective recognition performance was equivalent in the nap and wake groups. Collectively, these findings imply that the memory effects of sleep and TMR observed in this study were specific to category recall. Adjective recognition data is available in Table S1.

### Category Recall

#### Cue Frequency

In the nap group, category recall performance was superior for cued relative to non-cued items at T3. However, this behavioural benefit of TMR indexed as [cued category recall – non-cued category recall], was not predicted by the frequency of individual cue presentations in sleep (across all participants and cues, mean ± SD = 5.96 ± 5.85, *r*=.19, *P*=.34).

#### Objects vs. Scenes

We repeated the analyses reported in the main text to examine objects and scenes separately. A 2 (Type: Object/Scene) X 2 (Group: Nap/Wake) mixed ANOVA showed that object and scene recall did not differ at T1 (Type main effect: *F*(1,44)=0.23, *P*=.64), for either the sleep group or the wake group (Type*Group interaction: *F*(1,44)=1.54, *P*=.22). Again, there was no main effect of Group (*F*(1,44)=0.59, *P*=.45).

The factor ‘Type’ was added to our analysis of category retention at T3. A main effect of Type *F*(1,44)=6.88, *P*=.01) indicated that objects were generally better recalled than scenes. However, there was no interaction between Type and any other factor(s) (all *P*>.05), suggesting that the memory effects of sleep and TMR observed in this study did not vary according to category membership.

There was also a main effect of Type when this factor was included in our analyses of category retention at T2 (*F*(1,44)=4.11, *P*=.05). Again, there was no interaction between Type and any other factor(s) (all *P*>.05). Category recall data separated for objects and scenes is available in Table S2.

**Table S1:** Adjective recognition performance (%) at each test phase. The data refer to hits, misses, correct rejections (CR) and false alarms (FA). The sensitivity index (*d’*) was calculated for each test. Data are shown as means ± SEM.

|  |  | T1 |  |  |  |  |
| --- | --- | --- | --- | --- | --- | --- |
|  |  | Hit | Miss | CR | FA | d' |
| Nap |  | 84.04 | 15.96 | 96.37 | 3.63 | 2.89 |
|  |  | (± 2.10) | (± 2.10) | (± 0.69) | (± 0.69) | (± 0.10) |
| Wake |  | 89.74 | 10.26 | 96.42 | 3.58 | 3.15 |
|  |  | ± (1.71) | ± (1.71) | (± 0.77) | (± 0.77) | (± 0.14) |
|  |  | T2 |  |  |  |  |
|  |  | Hit | Miss | CR | FA | d' |
| Nap |  | 77.70 | 22.30 | 96.67 | 3.33 | 2.67 |
|  |  | (± 2.54) | (± 2.54) | (± 0.72) | (± 0.72) | (± 0.11) |
| Wake |  | 79.32 | 20.68 | 94.95 | 5.05 | 2.78 |
|  |  | (± 3.27) | (± 3.27) | (± 1.98) | (± 1.98) | (± 0.16) |
|  |  | T3 |  |  |  |  |
|  |  | Hit | Miss | CR | FA | d' |
| Nap |  | 79.74 | 20.26 | 96.52 | 3.48 | 2.83 |
|  |  | (± 2.86) | (± 2.86) | (± 0.87) | (± 0.87) | (± 0.10) |
| Wake |  | 80.11 | 19.89 | 94.95 | 5.05 | 2.78 |
|  |  | (± 3.38) | (± 3.38) | (± 2.46) | (± 2.46) | (± 0.11) |

### Discrimination Task

At the end of the experiment, participants were re-presented with all of the adjectives from the encoding phase and, for each, asked to indicate whether or not it was replayed in the offline period. The discrimination task analysis was restricted to items that were correctly recalled at T1 (and thus assigned to the cued and non-cued conditions). Cued stimuli that were and were not correctly identified as such were marked as hits and misses, respectively. Non-cued stimuli that were and were not correctly identified as such were marked as correct rejections or false alarms, respectively. A discrimination index was then calculated for each participant by subtracting the proportion of non-cued trails marked as false alarms from the proportion of cued trials marked as hits. The discrimination index was not significantly different from zero in either the nap group (*t*(26)=0.07, *P*=.95) or the wake group (*t*(18)=1.67, *P*=.11), implying that any memory effects of TMR were not due to conscious processing of the cues. It should be noted that while some of the wake group participants claimed to have heard the cues in the offline period, none of the sleep group participants professed any knowledge of adjective replay.

**Table S2:** Category recall performance (%) at each test phase for objects and scenes. Data are shown as means ± SEM. *Refers to the proportion of T1-recalled categories that were also recalled at T2. **Refers to the proportion of T2-recalled categories that were also recalled at T3.

| **Objects** | **T1** | **T2\*** |  | **T3\*\*** |  |
| --- | --- | --- | --- | --- | --- |
|  |  | *Not Cued* | *Cued* | *Not Cued* | *Cued* |
| *Nap* | 48.96<br>($\pm 2.92$) | 80.86<br>($\pm 2.36$) | 80.20<br>($\pm 3.16$) | 90.09<br>($\pm 2.03$) | 96.14<br>($\pm 1.76$) |
| *Wake* | 47.37<br>($\pm 3.51$) | 67.47<br>($\pm 4.51$) | 68.01<br>($\pm 4.32$) | 84.54<br>($\pm 4.02$) | 88.72<br>($\pm 3.55$) |
| **Scenes** | **T1** | **T2\*** |  | **T3\*\*** |  |
|  |  | *Not Cued* | *Cued* | *Not Cued* | *Cued* |
| *Nap* | 51.33<br>($\pm 3.18$) | 78.14<br>($\pm 2.70$) | 80.78<br>($\pm 2.69$) | 87.77<br>($\pm 1.85$) | 92.27<br>($\pm 1.50$) |
| *Wake* | 46.32<br>($\pm 2.77$) | 61.40<br>($\pm 4.57$) | 61.61<br>($\pm 4.53$) | 79.14<br>($\pm 4.90$) | 76.14<br>($\pm 5.62$) |

**Figure S1:**
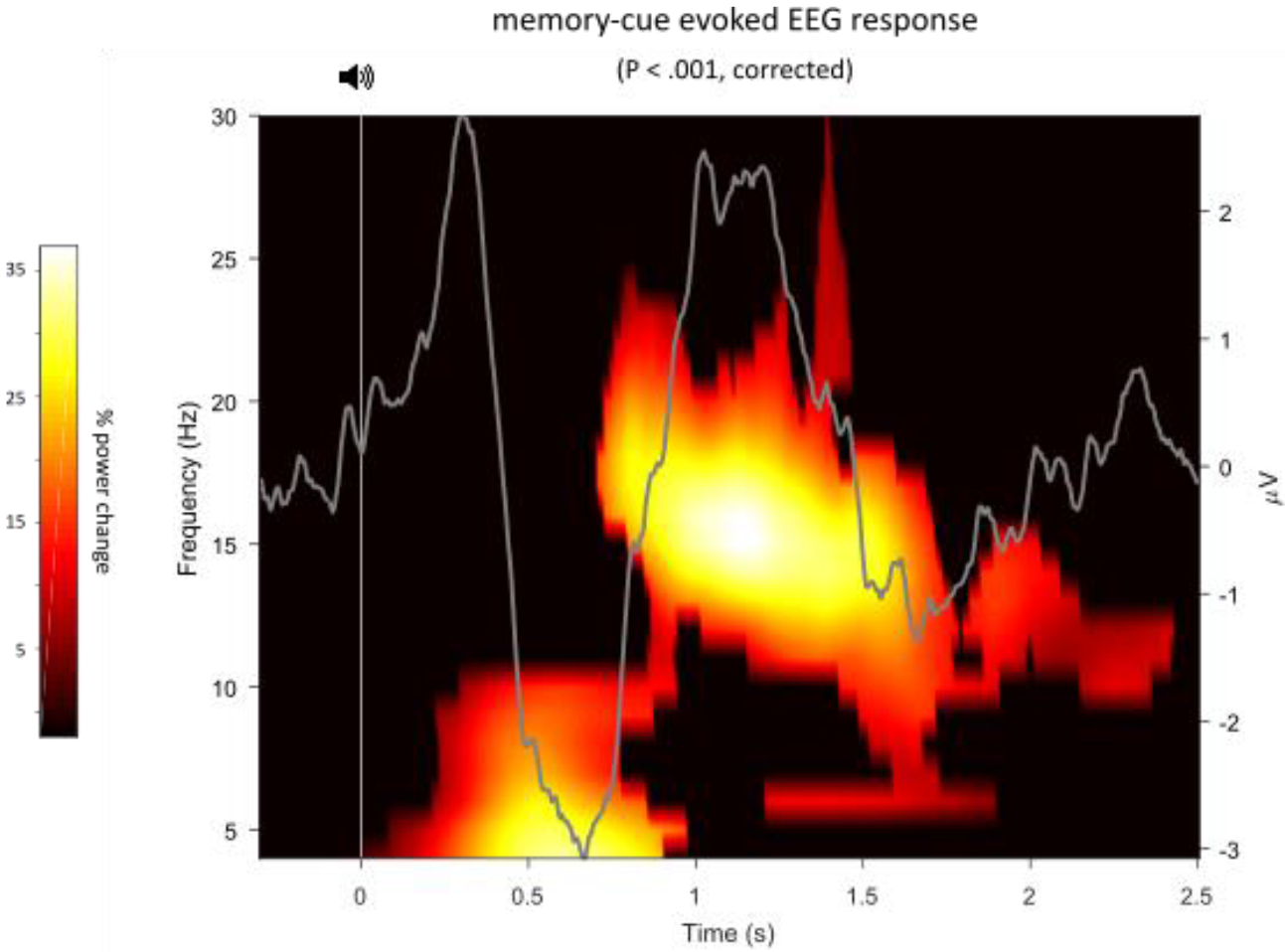
Significant increases in cue-evoked oscillatory power relative to a pre-cue baseline (−.3 to −.1 s). Results are thresholded at *P*< .001 (both for the initial, cluster-defining t test and for the final permutation test). Grey line reflects the corresponding ERP for old cues (cf. Figure 2B).

**Figure S2:**
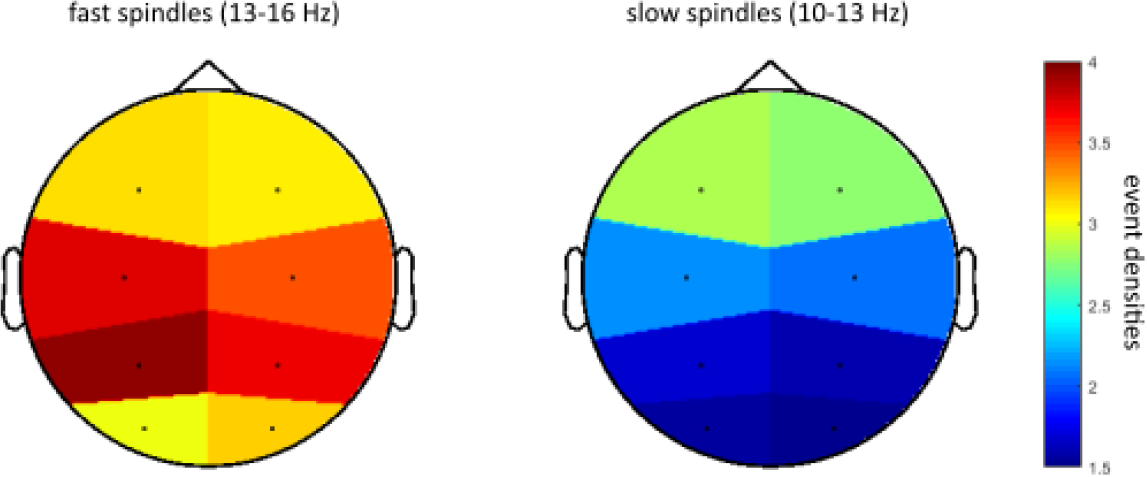
Event densities of fast (left; 13-16 Hz) and slow (right; 10-13 Hz) spindles detected algorithmically during NREM sleep (stages N2 and N3). Note the relative prevalence of fast spindles over centroparietal sites and that of slow spindles over frontocentral sites.

**Figure S3:**
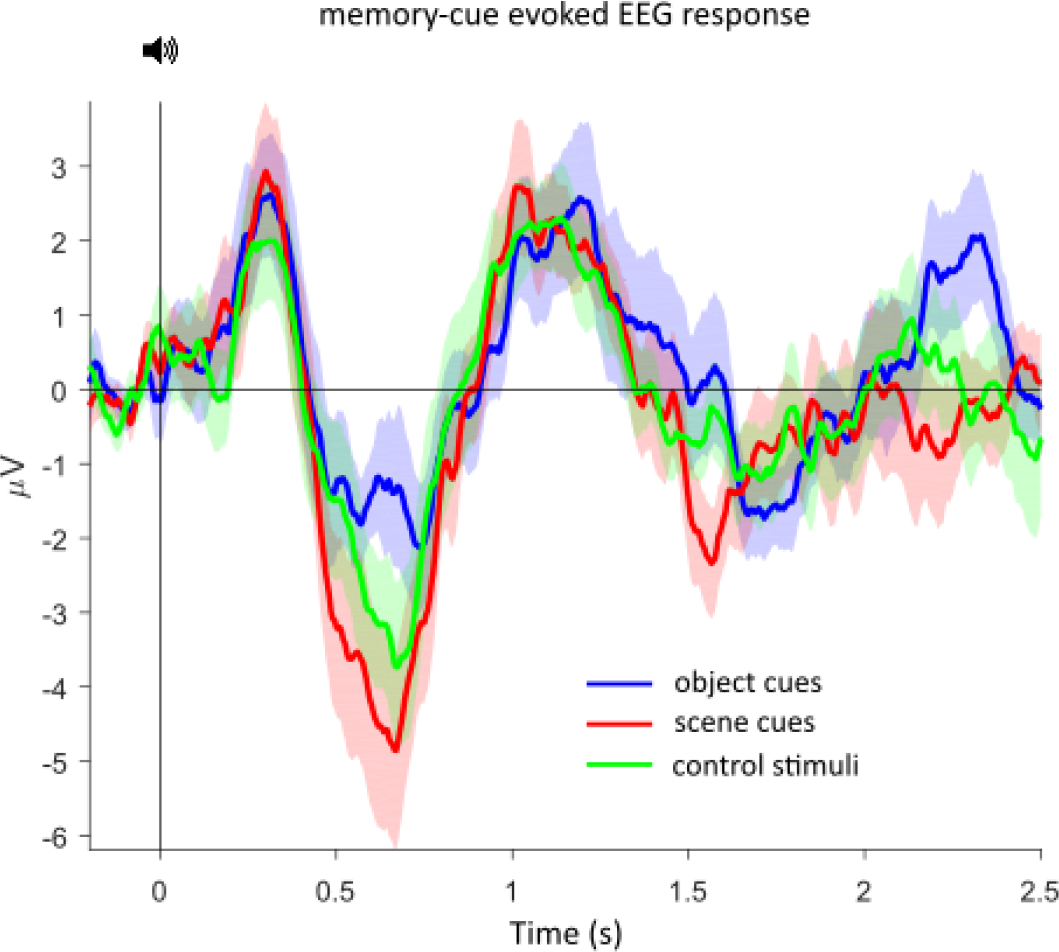
Event related potentials (ERPs) in response to adjective stimuli. Traces show mean ± SEM across participants, collapsed across all electrodes. A repeated-measures ANOVA revealed no significant condition effects comparing object cues, scene cues and control adjectives after controlling for multiple comparisons across time and electrodes.

**Figure S4:**
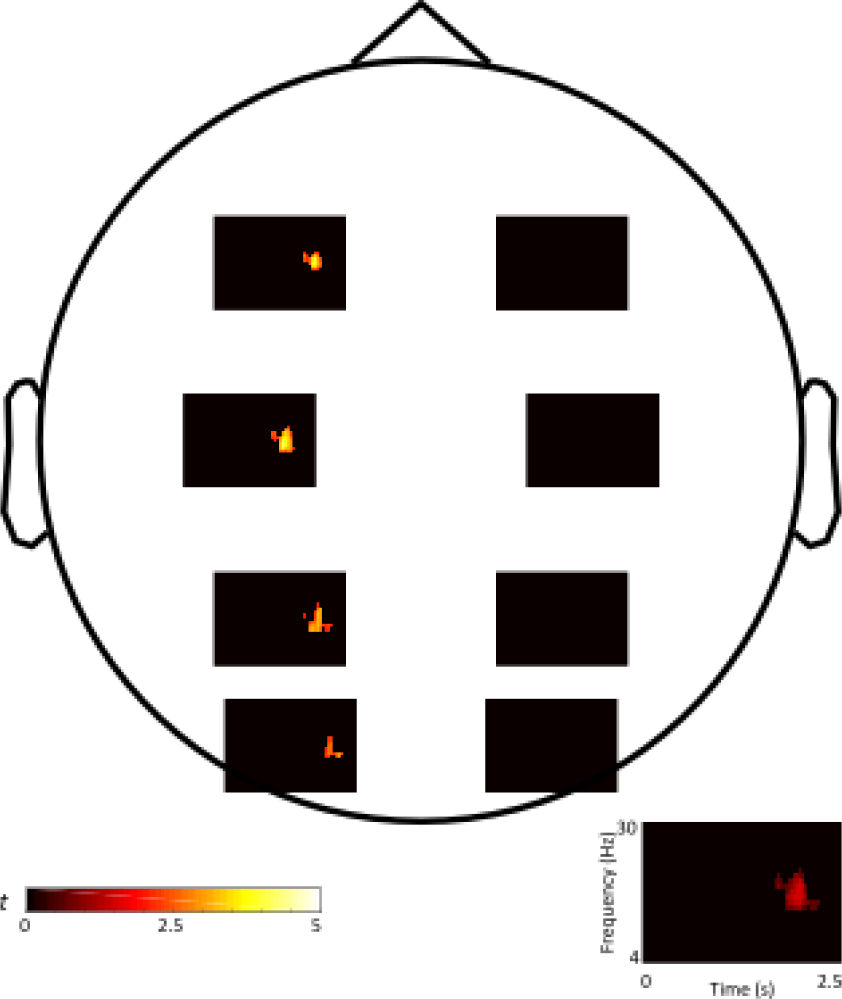
Topographical representation of significant clusters resulting from the contrast old TMR cues vs. novel control adjectives. Note the specificity of the effect in left hemisphere sites, with the maxima at C3 and P3. Right bottom: Effect integrated across all significant sites (cf. Figure 3B).

